# CRISPR/Cas-9 mediated knock-in by homology dependent repair in the West Nile Virus vector *Culex quinquefasciatus* Say

**DOI:** 10.1101/2021.01.14.426696

**Authors:** Deepak-Kumar Purusothaman, Lewis Shackleford, Michelle A. E. Anderson, Tim Harvey-Samuel, Luke Alphey

## Abstract

*Culex quinquefasciatus* Say is a brown, medium sized mosquito distributed widely in both tropical and subtropical regions of the world. It is a night-active, opportunistic blood-feeder and is responsible for vectoring many animal and human diseases, including West Nile Virus and avian malaria. Current vector control methods (e.g. physical / chemical) are increasingly ineffective; use of insecticides also imposes some hazards to both human and ecosystem health. Recent advances in genome editing have allowed the development of genetic methods of insect control, which is species-specific and, theoretically, highly effective. CRISPR/Cas9 is a bacteria-derived programmable gene editing tool that has been shown to be functional in a range of species. We demonstrate here, the first successful germline gene knock-in by homology dependent repair in *C. quinquefasciatus*. Using CRISPR/Cas9, we integrated exogenous sequence comprising a sgRNA expression cassette and marker gene encoding a fluorescent protein fluorophore (Hr5/IE1-DsRed, Cq7SK-sgRNA) into the kynurenine 3-monooxygenase (*kmo*) gene. We achieved a minimum transformation rate of 2.8% similar to rates achieved in other mosquito species. Precise knock-in at the intended locus was confirmed by sequencing. Insertion homozygotes displayed a white eye phenotype in early-mid stage larvae and a recessive lethal phenotype by pupation. This work shows an alternative and efficient method for genetic engineering of *C. quinquefasciatus*, providing a new tool for researchers interested in developing genetic control tools for this vector.

## Introduction

*Culex quinquefasciatus* Say also known as the southern house mosquito, is part of the *Culex pipiens* complex. The female mosquito is considered to be ornithophilic, establishing it as a major vector of many veterinary diseases, including avian malaria, which has been identified as a key factor in a number of extinctions of avian species, and a significant pressure on currently endangered ones^1^. However these opportunistic blood-feeders will also target mammalian hosts, bridging the gap of disease transmission from a range of different species from avian to mammalian hosts, thus posing a significant threat to human health, as seen with West Nile virus (WNV) ^2^. In the USA alone since the introduction of *C. quinquefasciatus* in 1999, there have been 48,000 human cases of WNV, for which the mosquito acts as an efficient vector^3^. Since humans are thought to be a dead-end host for WNV, each of these likely represents mosquito-vectored avian-to-human transmission. Horses are also vulnerable to WNV, with about 25,000 cases and a case fatality rate of about 33%^4^. In addition, it is a competent vector of the St. Louis encephalitis virus and eastern equine encephalitis virus^5^.

The major human health impact of *C. quinquefasciatus* globally is as a vector of lymphatic filariasis (LF). LF is a neglected tropical disease caused by a nematode, which can live up to 6-8 years inside their human host, causing disruption and permanent damage to the lymphatic system; despite extensive control efforts there are still an estimated 50 million cases worldwide^6,7^. Eradication of this disease is presently unexpected despite great efforts with mass drug administration programs, leading to increasing focus on complementary vector control strategies^8^.

Vector control methods currently used for other mosquito species such as removing or chemically treating larval habitats are limited to their effectiveness in large scale implementation. Moreover, some breeding sites may be impractical or difficult to locate^9^. In addition, most current control strategies are heavily dependent on the usage of insecticides, which can have major impacts on non-target species supporting local ecosystems^10^. Furthermore, insecticide usage is under threat of becoming ineffective in the target population due to resistance either by mutations in an individual target site gene, or by a rise in metabolic resistance, which essentially allows the insect to degrade the active insecticide more rapidly^11-13^

New strategies and targets for vector control are therefore urgently required. Genetic approaches potentially provide this, and may be more suitable for large scale implementation, in addition to having a lesser impact on non-target species^10,14^.The ability to edit genes allows the characterization of new targets, but also opens the door for the implementation of genetic manipulation of key vector populations. This might aim to suppress the vector populations, or insert genes which reduce vector competence. Introgressing such traits into wild vector populations might be through mass-release, or by using gene drive systems to amplify the effect of relatively small initial releases^14,15^. The recent availability of efficient gene-editing tools such as clustered regulatory interspaced short palindromic repeats-associated protein 9 (CRISPR/Cas9) has made many of these approaches more feasible. The Cas9 endonuclease is typically paired with a synthetic single guide RNA (sgRNA); the sgRNA sequence will have complementary bases to that of the target site region of DNA in the genome. When the sgRNA and DNA bind, the Cas9 protein induces a double stranded break in the DNA. This can be repaired either by non-homologous end joining (NHEJ) or homology directed repair (HDR)^16^. Repair through the NHEJ pathway typically results in small indel (insertion/deletion) mutations which can be useful in assessing the function of target genes, if they result in a frame-shift or loss of an important protein function (knock-out). HDR-based repair can be utilised to ‘knock-in’ an exogenous DNA sequence, if such a DNA sequence is provided with flanking regions homologous to the endogenous break site, for example by co-injection of a ‘repair-template’ plasmid alongside CRISPR components. Alternatively, sgRNA and Cas9 components can be integrated into the germline of a target species whereby the two repair pathways can be harnessed as mechanisms for gene drive. For example, HDR can be used for homing-based drives^17-19^ while NHEJ repair of essential genes can form the basis for ‘break and repair’ based systems such as CLVR or TARE^20,21^.

Regarding mosquitoes, CRISPR/Cas9 has been used successfully to generate germline and somatic knock-in mutations in *Aedes* and *Anopheles* species’ through the HDR repair mechanism^17,22-26^. However, to date, only use of the NHEJ pathway to generate ‘knock-out’ mutations has been reported in *C. quinquefasciatus*^27-30^. Only a few previous studies have successfully generated transgenics using transposon-mediated transformation in this species, using a *Hermes*-based vector^31,32^. Our own attempts to generate transgenics with *piggyBac*-based vectors did not recover any transgenics using both plasmid^33^ and *in vitro* transcribed mRNA^34^ as transposase sources.

In this study, using the kynurenine 3-monooxygenase (*kmo*, also known as kynurenine hydroxylase or *kh*) gene as a target, we demonstrate for the first time the ability of the CRISPR/Cas9 system to generate knock-in mutations in *C. quinquefasciatus*. As well as a successful proof of concept for this technology, our chosen integrated components (an RNA Polymerase III promoter expressing a sgRNA, itself targeting the knock-in site) is the first step in assessing the potential of a CRISPR/Cas9 homing drive-based approach in this globally important mosquito pest.

## Results

### CRISPR/Cas9 based site specific insertion into *kmo*

Our earlier work identified a region homologous to the *Aedes aegypti kmo* gene which yielded a white-eyed phenotype when disrupted by CRISPR/Cas9^27^. The target site of the most active sgRNA (LA935) was selected for knock-in experiments. Two independent rounds of embryonic microinjections were performed using a CRISPR/Cas9 HDR donor template (Fig 1). In the initial round of injections 384 embryos were injected. Out of these 170 hatched as first instar larvae of which 57 (14.8%) survived to adulthood. In total 7 pools were generated from these injection survivors and four ovipositions of G_1_ larvae were collected and screened (Table 1). No fluorescent larvae were identified from this series of injections, from a total of 5,313 G_1_ larvae screened. A subsequent series of injections were performed on 601 embryos and of these 141 larvae hatched and 71 survived to adulthood (11.8%). In total 4,441 G_1_ larvae were screened and we identified larvae with DsRed fluorescence in two pools. A total of 60 positive individuals were identified from pool K and 48 from pool L. The HDR integration rate for the second round of injections was calculated to be a minimum of 2.8%; combining both experiments would indicate a minimum transformation rate of 1.6%. “Transformation rate” is typically defined as the number of independent integration events identified per fertile G_0_ adult injection survivor. Pooling G_0_, required here for efficient recovery of G_1_, means that we do not know what proportion of G_0_ adults were fertile; the efficiency calculation therefore uses total G_0_ adults instead. The calculated rate also assumes that all positive G_1_ larvae recovered from a single pool were the result of a single integration event, here assuming that the 60 positives identified represent two integration events; if this were not the case then this calculation is an underestimate of the true integration rate. For site-specific integration, independent insertions within one pool are not readily distinguished.

**Table 1.** Results of two sets of injection experiments aiming to integrate a knock-in cassette into the *kmo* locus. *Culex wild-type* embryos co-injected with AGG2069 HDR donor plasmid, LA935 sgRNA and spCas9 protein. Adult injection survivors were mated in the number of pools indicated. These were blood fed and four ovipositions collected and larvae screened for DsRed fluorescence. Integration rate is calculated by dividing the number of pools which yield an integration event (here 2) by the total number of G_0_ adults.

| Experiment | No. injected | G<sub>0</sub> adult survivors (%) | G<sub>0</sub> pools | G<sub>1</sub> screened | DsRed positive | Minimum integration rate |
| --- | --- | --- | --- | --- | --- | --- |
| 1 | 384 | 57 (14.8%) | 7 | 5313 | 0 | 0 |
| 2 | 601 | 71 (11.8%) | 7 | 4441 | 60 (pool K)<br>48 (pool L) | 2.8% |
| Combined | 985 | 128 (13%) | 14 | 9754 | 108 (2 pools) | 1.6% |

**Figure 1.**
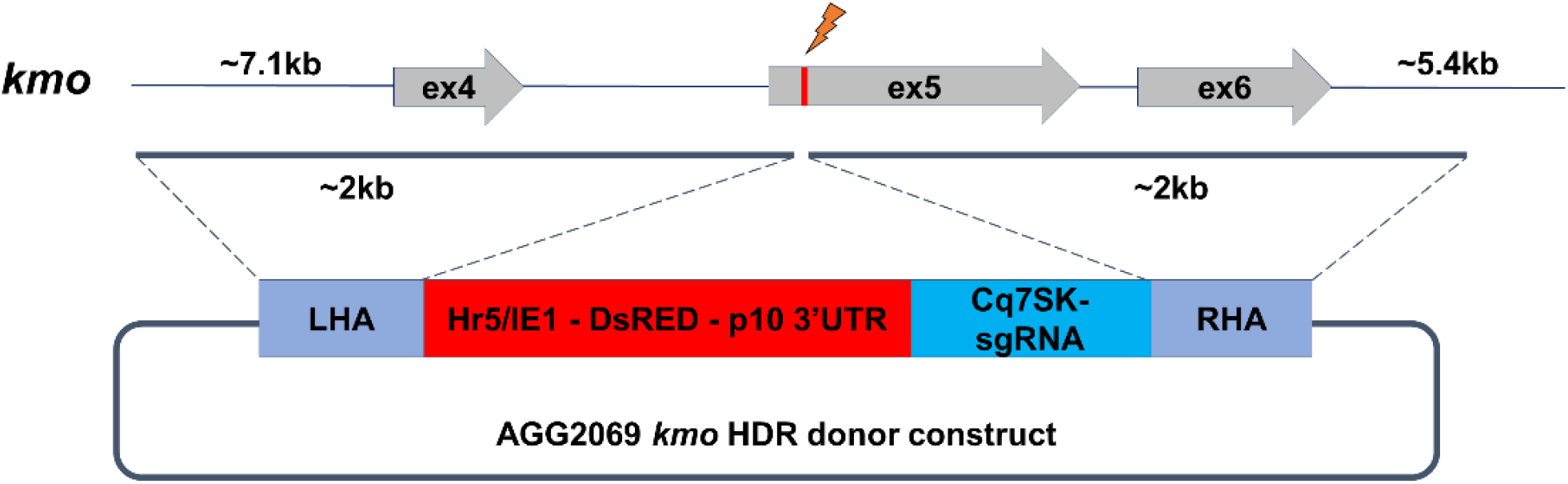
CRISPR/Cas9 based *kmo* knock-in cassette. Representation of the *kmo* locus and HDR donor construct for integration. Grey arrows represent exons and the red line indicates the sgRNA target site within exon 5. The blue lines indicate the left and right homology arm sequences.

### Molecular confirmation of *kmo* insertion

Successful knock-in of the HDR construct at the *kmo* locus was confirmed by PCR (Fig 2). Representative fluorescent individuals from both pools K and L produced amplicons of expected size. This PCR assay produced some multiple non-specific amplification products in WT samples, so putative integrations were confirmed by Sanger sequencing (Fig S2). A second diagnostic PCR with the two external (genomic) primers (LA6087 & LA6088) produced an amplicon of expected size (∼4kb) in the WT DNA sample and two amplicons (∼ 4kb & ∼ 7kb) in the AGG2069 pool K & L samples confirming the HDR insertion in one of the WT *kmo* alleles (Fig S1).

**Figure 2.**
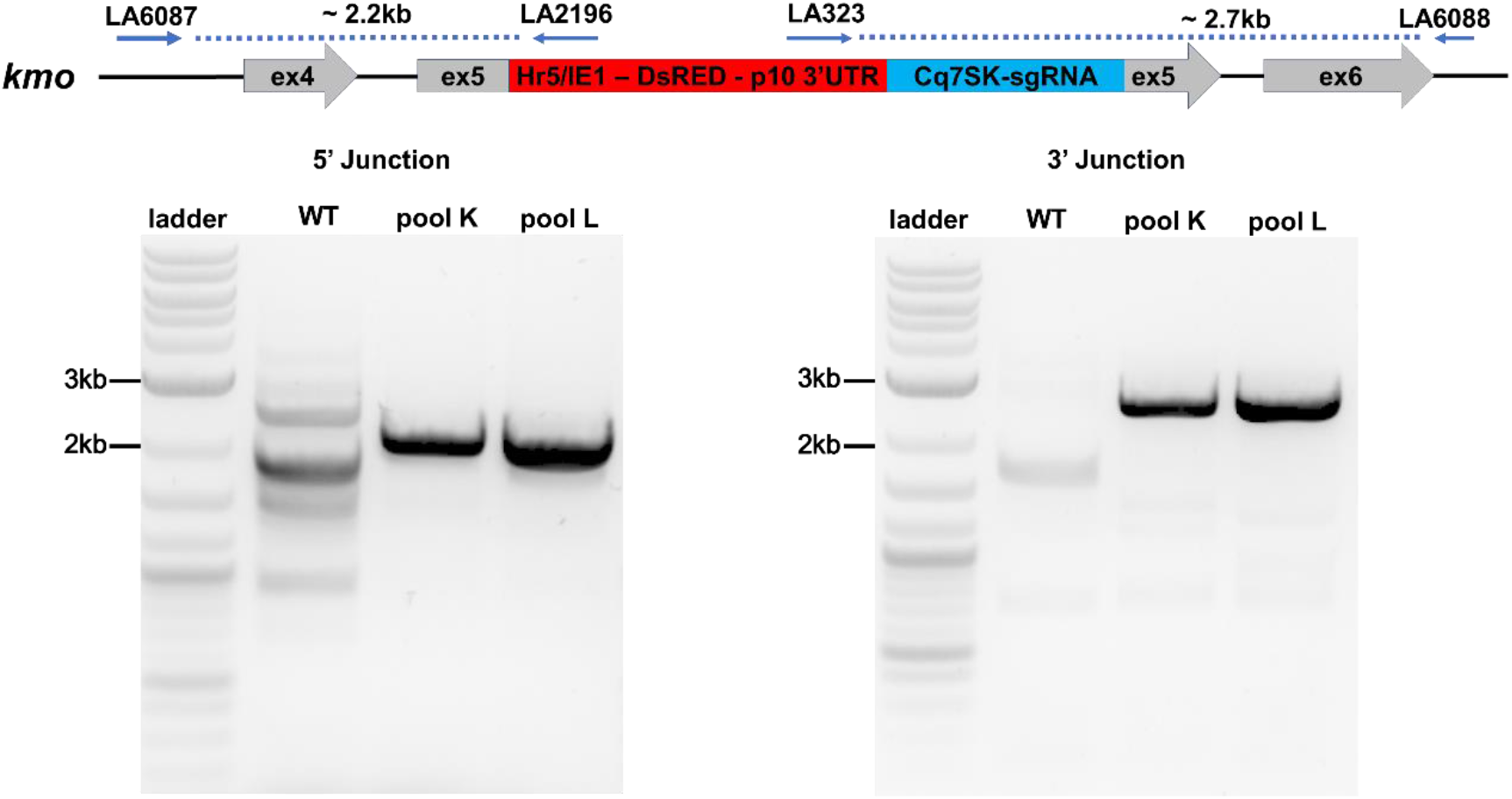
PCR confirmation of the integration of the HDR cassette at the *kmo* locus. The blue arrows on the diagram indicate the primers used for PCR confirmation. Primers LA6087 and LA6088 were selected to anneal to sequences outside the homology arms of AGG2069 (Fig 1). The blue dotted lines show the PCR amplicon. DNA ladder in the first lane is the NEB Quick-Load Purple 1 kb Plus DNA ladder.

### Generation of homozygous lines

To assess the viability of homozygous *kmo* insertions, G_1_ fluorescent individuals from both pools K and L were sibling crossed. The inheritance pattern in the offspring (G_2_) was expected to be 25% homozygous for *kmo* knock-in (red fluorescent and white-eyed), 50% heterozygous (red fluorescent and WT eyed) and 25% wildtype (non-fluorescent, WT eyed). The phenotypes of individuals of these three genotypes are shown in Fig 3. Larvae which were homozygous for the *kmo* knock-in displayed brighter fluorescence when compared to the heterozygotes (Fig 3a), presumably due to having two copies of the transgene. Loss of eye pigmentation (white-eyed phenotype) and bright fluorescence was observed in 19.9% of the G_2_ larvae in pool K and 15.8% in pool L (Table 2) identifying these individuals as homozygous for the insertion. Heterozygotes (56% in pool K and 51.5% in pool L) and wild-type (24.1% in pool K and 32.7% in pool L) were observed at approximately mendelian rates (Table 2). The observation of white eyes only when associated with bright, fluorescent individuals provides additional evidence to suggest that the HDR integration sites are within the *kmo* locus. The decrease from the expected 25% frequency, as well as an observed slow growth of homozygous individuals suggests that high fitness costs are associated with these insertions. Supporting this conclusion, we observed that no homozygous transgenic larvae survived to pupation, while wild-type and heterozygous larvae pupated normally.

**Table 2.** Genotype frequencies arising from sibling crosses. Heterozygotes from pools K and L were sibling crossed and their offspring screened for inheritance of the transgene as well as eye color phenotype as late larvae.

| Pool | No. of ovipositions screened | Homozygotes (white eyes, DsRed +) larvae | Heterozygotes (dark eyes, DsRed +) larvae | Wild-type (dark eyes, DsRed -) larvae | Total larvae screened |
| --- | --- | --- | --- | --- | --- |
| K | 1 | 38 (19.9%) | 107 (56%) | 46 (24.1%) | 191 |
| L | 2 | 43 (15.8%) | 140 (51.5%) | 89 (32.7%) | 272 |

**Figure 3.**
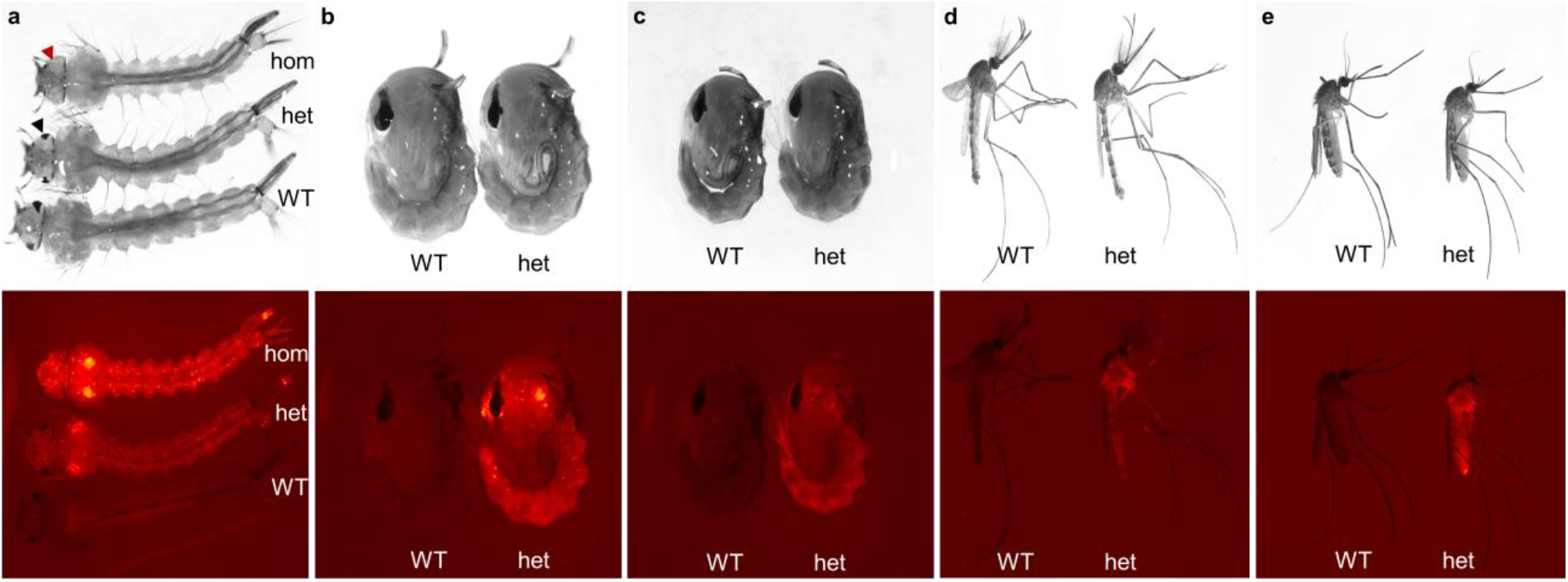
Phenotype of *kmo* knock-in in different life stages of *C. quinquefasciatus* (AGG2069 and wild-type). a) Photomicrographs of late larvae: white light (top) and red fluorescence (bottom). Pigmented eyes are clearly visible in wildtype (WT) and heterozygotes (het, one eye is indicated with a black arrowhead); this eye pigment is missing in homozygotes (hom, one eye indicated with red arrowhead). b) Male pupae under white light (top) and red fluorescence (bottom). c) Female pupae under white light (top) and red fluorescence (bottom). d) Male adults under white light (top) and red fluorescence (bottom). e) Female adults under white light (top) and red fluorescence (bottom).

## Discussion

Development of next-generation genetics-based control strategies such as gene drives require an efficient and precise method for integrating transgenic sequences into the germline of a target organism. Previous efforts to utilise the *piggyBac* transposase system for this purpose in *C. quinquefasciatus* were unsuccessful, despite multiple different forms of transposase being utilised^31^. This was surprising, given the extremely broad range of species, across multiple phyla, in which *piggyBac* has been shown to be an efficient tool for germline integration^35^. The reasons for this curious lack of success in *C. quinquefasciatus* are yet to be determined but could, for example, be due to a susceptibility of the *piggyBac* mRNA sequence to endogenous silencing pathway components such as PIWI or other small silencing RNAs. Alternatively, the stability of the *piggyBac* protein, or its ability to interact with DNA may be disrupted in *C. quinquefasciatus*. Whilst this poses interesting fundamental questions, the potential damage caused by this mosquito species, and its rapid spread into new habitats around the globe, necessitates the rapid development of orthogonal technologies to act as the building blocks for novel control systems such as gene drives. Previous work demonstrated that the *Hermes* transposase system was functional in *C. quinquefasciatus*, however, resulted in the non-canonical integration of plasmid backbone sequences alongside those ‘desired’ components within the *Hermes* flanks. This is undesirable for the development of genetic control tools designed to be released into the wild as the presence of these sequences would likely complicate the regulatory process for such lines. As with the *piggyBac* system, *Hermes* also operates through a semi-random integration method, making it inappropriate for many of the more powerful gene drive designs, which require precise integration of transgenic components into target loci.

Here we provide a solution to both these issues by demonstrating the functionality of CRISPR/Cas9-based knock-in in *C. quinquefasciatus*. Assessment of the flanking regions of the two established lines showed that integration was precise at the target cut site with no integration of undesirable backbone components. Additionally, our 1.6% transformation rate (2.8% for the second round of injections) suggests a relatively efficient process and certainly one which is efficient enough for the testing of different exogenous components. As an example, our knock-in cassette was built to include an RNA Polymerase III promoter previously found to be highly active in *C. quinquefasciatus* Hsu cells^36^ which drove *in vivo* expression of the same sgRNA used to integrate the cassette. This represents the first step towards testing of the homing-drive concept in *C. quinquefasciatus* through a ‘split-drive’ design^19^. Further work will be required to assess the functionality of the chosen 7SK promoter to express the sgRNA in suitable germline tissues and to develop other lines capable of expressing Cas9 with compatible spatial and temporal characteristics.

Interestingly, during our experiments we observed a severe recessive fitness cost associated with the *kmo* transgene integration, resulting in death of all homozygous individuals prior to pupation. This was surprising for two reasons, the first is that our previous work generating a frame-shift deletion at this locus in *C. quinquefasciatus* using the same sgRNA as utilised to specify the integration site here did not result in such a lethal phenotype, although a significant sub-lethal fitness cost was observed^27^. Secondly, our unpublished work generating a similar knock-in in the homologous exon of the *kmo* gene of another culicine mosquito, *Aedes aegypti*, did not result in a recessive lethal phenotype. The situation in *C. quinquefasciatus* appears to share more in common with the relatively distantly related *Anopheles stephensi*, where severe fitness costs associated with *kmo* knock-in / *kmo* knockout homozygotes resulted in high levels of adult female lethality post blood-feeding, and significant reductions in egg-laying for surviving females^24^ – though little apparent effect on males, whereas lethality affected both sexes in our *C. quinquefasciatus* knock-in lines. This fitness cost was harnessed as a resistance management mechanism in next-generation *A. stephensi* gene drives, providing a framework for similar designs in *C. quinquefasciatus*^*37*^. The basis for the observed differences between homozygous viability for our *C. quinquefasciatus* knock-in and knock-out lines is unclear. It may be that the original mutant lines retain some activity, i.e. are hypomorphic, even though sequencing identified them as frame-shift mutants – possibly there are alternative splicing variants, or a truncated protein retains some activity. It is also possible that the observed lethality is associated with a closely linked background mutation, though it seems unlikely that this would be present in both knock-ins but not the original knock-outs, all of which were generated from the same wild-type colony. Further research is required to explore this and other potential explanations.

It is hoped that this work will provide a springboard for those researchers interested in developing homing-based and other gene drive strategies in this pernicious global pest.

## Materials and methods

### Mosquito rearing

Both wild type TPRI (Tropical Pesticides Research Institute) *C. quinquefasciatus*^27^ and the *kmo* HDR knock-in lines (AGG2069) were maintained at 28°C, 70% humidity and 12 hr day-night cycle in an insectary as previously described^27^. Egg rafts were collected from adult cages in a 150 ml plastic container filled with horse hay infused water. Mosquito larvae were fed with pelleted pond fish food. Adult mosquitoes were fed *ad libitum* with 10% sucrose solution. A Hemotek system (Hemotek, Blackburn, UK) was used to provide debrifinated horse blood (TCS Biosciences, Buckingham, UK) through sausage casing and a layer of Parafilm.

### Plasmid design and cloning

The kynurenine 3-monooxygenase (*kmo*) gene (CPIJ07147) of *C. quinquefasciatus* was previously identified and sequence confirmed as described^27^. Approximately 2 kb upstream and downstream of the precise sgRNA cut site was used as homology arms. A cassette containing the Cq7SK promoter^36^ was PCR amplified from the plasmid AGG1127 and the sgRNA sequence was added in using the oligos designed for the PCR. The Hr5/IE1-DsRed-p10 3’UTR cassette was amplified by PCR from another plasmid (AGG1906). All the fragments were gel purified and assembled using the NEBuilder HiFi DNA Assembly Master Mix (New England Biolabs, Ipswich, MA, USA). Complete AGG2069 plasmid sequence has been deposited to NCBI (Accession number: MW417419).

### CRISPR/Cas9 injection mix components

Purified NLS-SpCas9 protein was procured from PNA Biosciences (Thousand Oaks, CA USA). Nuclease free DEPC treated water was used to resuspend the lyophilized Cas9 protein to a concentration of 1500 ng/µl and stored at −80°C. sgRNA LA935^27^ was generated from a PCR DNA template (overlapping primers LA935 5’-GAAATTAATACGACTCACTATAGGACAGTGCGGTCCG-CAAGGGTTTTAGAGCTAGAAA-3’ and LA137 5’-AAAAGCACCGACTCGGTGCCA-CTTTTTCAAGTTGATAACGGACTAGCCTTATTTTAACTTGCTATTTCTAGCTCTAAAAC-3’) containing the T7 promoter using the MEGAscript T7 Transcription kit (ThermoFisher Scientific, Walthum, MA USA). The reaction mix was incubated at 37°C for 16 hours and purified using the MEGAclear Transcription Clean-Up kit (ThermoFisher Scientific, Walthum, MA USA). Injection mix (20 µl) was made with the following components: AGG2069 (HDR donor plasmid, 800 ng/µl), LA935 sgRNA (40 ng/µl), Cas9 protein (300 ng/µl), 10X Injection Buffer^38^ (2 µl), DEPC water (up to 20 µl). The assembled injection mix was incubated at 37°C for 20 minutes to pre-complex the Cas9 and sgRNA, then centrifuged at max speed and 4°C for at least 10 minutes. Mix was maintained on ice throughout injection.

### Embryonic microinjections

*Culex* embryo microinjections were performed as previously described^27^. A microscopic injection station equipped with FemtoJet 4X microinjector (Eppendorf, Hamburg, DE) was used for injections. Injections were carried out using quartz capillaries (0.7 mm internal diameter and 1.0 mm external diameter) pulled into needles using a Sutter P2000 laser based micro-pipette needle puller (Sutter Instruments, Novato, CA USA) and the following program: HEAT=729, FIL=4, VEL=40, DEL = 128, PUL = 134, Line =1.

A clear plastic cup containing approximately 100 ml of hay infused water was placed into the adult cages on the 5^th^ day after a blood meal. Cages were placed in the dark to encourage egg laying and allowed to lay for 45-60 minutes. The egg rafts were disaggregated and aligned horizontally on a piece of moistened chromatography paper and against a nitrocellulose membrane (GE Healthcare, Amersham UK). Lines of embryos were then transferred to a piece of Scotch double-sided tape 665 (3M, USA) on a plastic coverslip. Prepared eggs were covered with Halocarbon oil 27 (Sigma Aldrich, Gillingham UK) to prevent desiccation and injected. Injected eggs were washed with distilled water to remove as much oil as possible and (still on the coverslip) submerged egg side down into larval rearing trays and allowed to hatch. Surviving larvae were transferred to a new tray with hay infused water and maintained at the standard rearing conditions.

### Crosses and screening

Both male and female adult injection survivors (G_0_) were mated to the parental wildtype strain (TPRI). Male G_0_ individuals were crossed to 3 wild type females, and after 7 days these were pooled into groups of 10 males. G_0_ females were mated in pools of 10 to approximately 20 wildtype males. Pools were blood fed and after 5 days, eggs were collected as described above. Four ovipositions were collected for all pools and screened under the Leica MZ165FC microscope (Leica Biosystems, Milton-Keynes UK) and images were taken using a Leica DFC camera and the settings: Brightness 82%, saturation 0 and Gamma 0.71 (for white light images) and Brightness 82%, Saturation 0.146 and Gamma 0.40 (mCherry - red fluorescence). Fluorescent marker positive G_1_ males and females were crossed to generate homozygous lines.

### PCR confirmation of HDR insertions

Genomic DNA from wildtype and DsRed individuals was extracted using the NucleoSpin Tissue kit (Macherey Nagel, Düren, Germany). The 5’ and 3’ junctions of the kmo integration were PCR confirmed using two internal primers LA2196 (5’-CCAGTTCGGTTATGAGCCGT-3’) and LA323 (5’-ACCAAATCTGCCAGCGTCAATAG-3’), that bind within the inserted cassette and two external primers LA6087 (5’-TTCGGTTTGCCCAAAGAAGC-3’) and LA6088 (5’-AAATGTTCGTCTCCGACCCC-3’) that bind to the genome external to the homology arms. An additional PCR was also performed with only the external primer set: LA6087 and LA6088 (Fig S1). Q5 High-fidelity 2X Master Mix (New England Biolabs, Ipswich, MA, USA) was used with the following cycling conditions: initial denaturation 98°C for 30 sec, 35 cycles of denaturation 98°C for 10 sec, annealing temperature 67°C for 10 sec and extension 72°C for 4 mins, final extension 72°C for 10 min, followed by hold at 4°C. The PCR amplicons were electrophoresed in a 1% agarose gel with SYBR Safe (ThermoFisher Scientific, Waltham, MA, USA). The amplicons of expected size were excised, gel purified using NucleoSpin Gel and PCR Clean-up kit (Macherey Nagel, Düren, Germany) and subjected for Sanger sequencing.

## Acknowledgements

LA and THS are supported through strategic funding from the UK Biotechnology and Biological Sciences Research Council (BBSRC) to The Pirbright Institute (BBS/E/I/00007033, BBS/E/I/00007038 and BBS/E/I/00007039). DP is funded by a PhD Studentship from The Pirbright Institute. LS and MAEA are funded by the Bill and Melinda Gates Foundation (INV-008549).

## Author contributions

THS, MAEA, and LA designed the experiments. LS and DP performed the experiments and analysed the data. All authors wrote, revised, and approved the manuscript.

## Competing interests

The authors declare no competing interests.

## Supplementary Information

**Figure S1.**
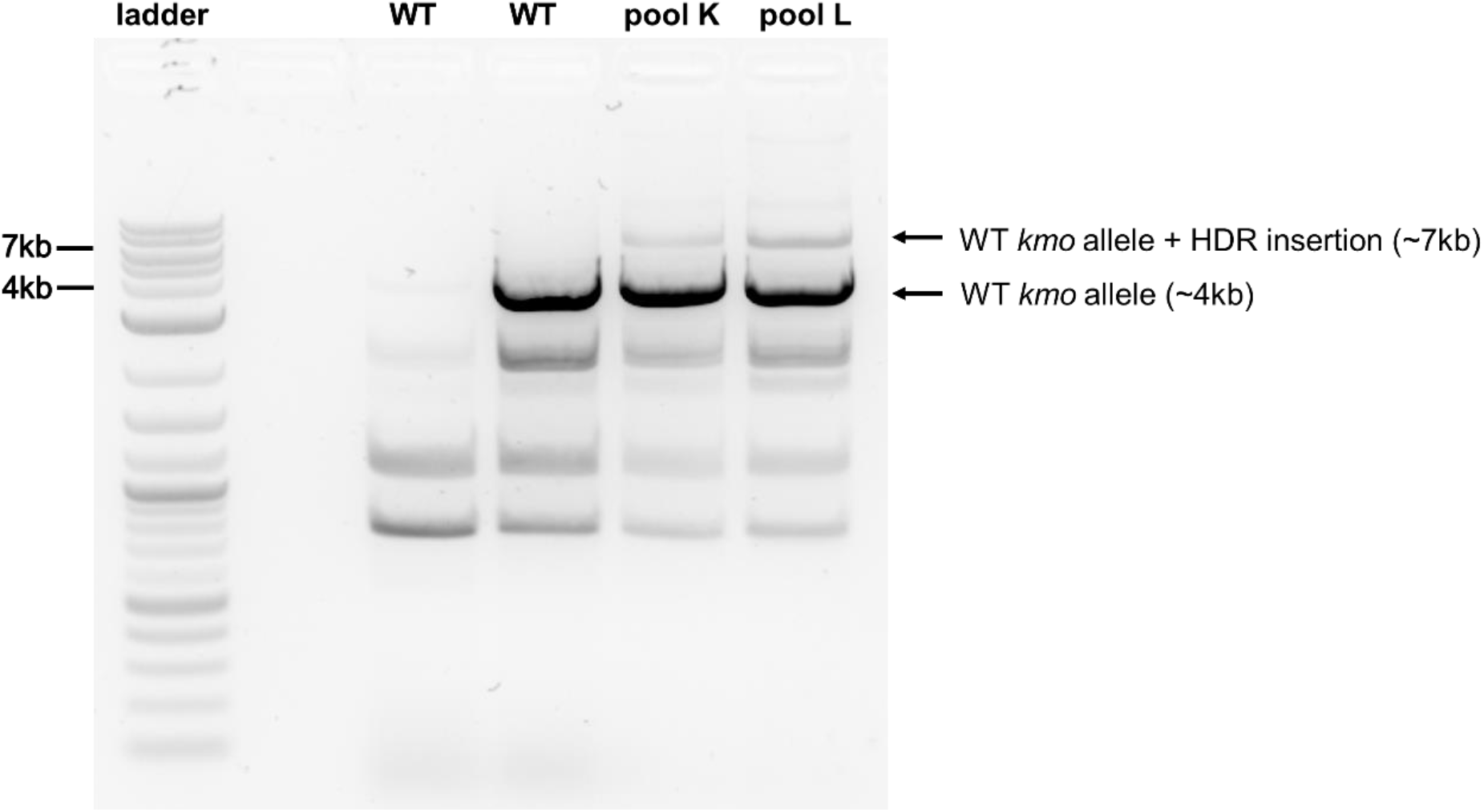
PCR confirmation of the *kmo* knock-in. Successful integration of the HDR construct into the *kmo* locus produced expected banding patterns along with additional seemingly non-specific amplicons. The amplicons of expected size for the *kmo* alleles were excised and sequence confirmed.

**Figure S2.**
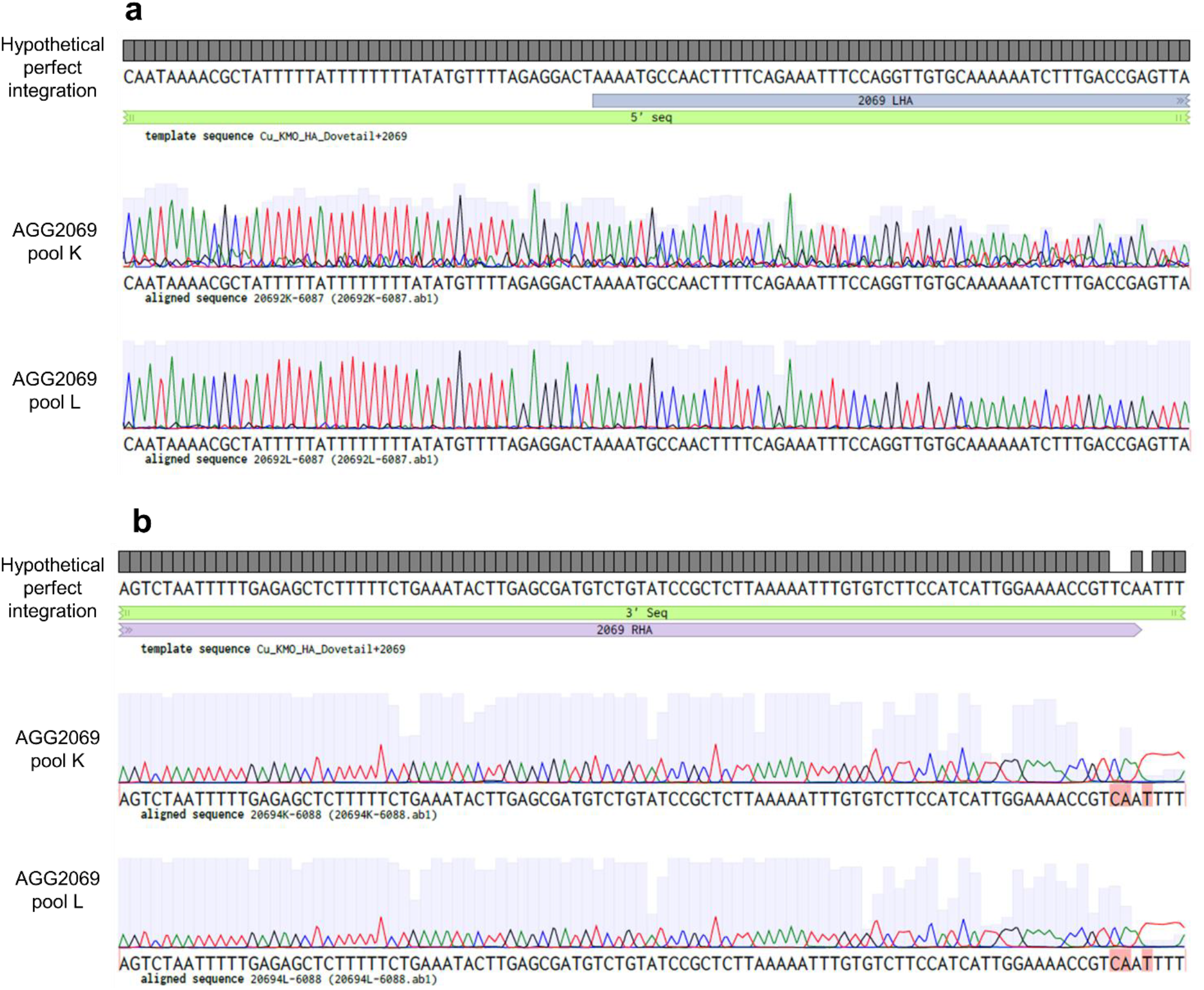
Sequence confirmation of the *kmo* knock-in. Sanger sequencing traces showing (a) the 5’ and (b) the 3’ junctions of the *kmo* knock-in cassette within the genome. In the 3’ junction, the reverse primer binds to the genomic region immediately adjacent to the right homology arm which resulted in poor quality basecalling (red highlighted bases) at the junction of the homology arm with the genome (b).

## Notes

### Competing Interest Statement

The authors have declared no competing interest.

